# Unsupervised single-cell clustering with Asymmetric Within-Sample Transformation and per cluster supervised features selection

**DOI:** 10.1101/2023.05.17.541148

**Authors:** Stefano Maria Pagnotta

## Abstract

This chapter shows applying the Asymmetric Within-Sample Transformation [14] to single-cell RNA-Seq data matched with a previous dropout imputation. The asymmetric transformation is a special winsorization that flattens low-expressed intensities and preserves highly expressed gene levels. Before a standard hierarchical clustering algorithm, an intermediate step removes non-informative genes according to a threshold applied to a per-gene entropy estimate.

Following the clustering, a time-intensive algorithm is shown to uncover the molecular features associated with each cluster. This step implements a resampling algorithm to generate a random baseline to measure up/down-regulated significant genes. To this aim, we adopt a GLM model [10] as implemented in DESeq2 [9] package.

We render the results in graphical mode. While the tools are standard heat maps, we introduce some data scaling so that the results’ reliability is crystal clear.

## 1 Introduction

Clustering addresses the question of aggregating items according to some metric of similarity. In bioinformatics, a general task is to cluster bulk samples or single cells, in both cases described by a large set of omic features such as transcriptomic expression levels. In this chapter, we show how to implement the AWST protocol [14] to single cells and get a clustering. The core of the protocol is an asymmetric data transformation that down-weights noisy features so that their contribution to the analysis vanish. In general any clustering procedure can follow AWST, but we, as long as data dimensionality allows, prefer hierarchical algorithms matched with the Ward’s linkage applied to an Euclidean distance matrix. Consistently with these choices, the criterion for exploring the number of clusters is the Calinski-Harabasz [3] curve. These statistical methodologies shares the key concept of maximizing variability between groups while minimizing the variability within. Once the cells partition has been computed, a downstream procedure to characterize every single cluster is definite.

Essentially the procedure performs a gene-wise ANOVA analysis with the help of the generalized linear models (GLM) as implemented in the DESeq2 package [10]. The large amount of variability induced in the data processed by the GLM assures robust results.

Although we present the two analyses in sequence (clustering + per cluster features selection), they can be separated to be included in other pipelines. Any clustering procedure can feed the group signature mining step. As well as any signature mining step can follow the proposed clustering procedure.

A few modifications also allow researchers to apply our proposal to bulk RNA-Seq data.

The chapter has been arranged as follows: Section one shows the results generated by the pipeline, with some molecular remark; section two shows materials and software; the last section describe the methodology step by step. A few notes close the chapter.

The software can be retrieved from [12].

## 2 Result

We consider the collection of 6,964 single cells from human breast epithelial tissue of donor [6] analyzed in [11]. Raw counts have been first treated for dropout imputation with [5] methodology, then the AWST pipeline [14] has been applied. We performed an unsupervised hierarchical clustering with Ward’s linkage and Euclidean distance using 2,141 features out of about 20,000 gene expression levels. The Calinski-Harabasz [3] method suggests the number of clusters and proposes to investigate first a partition of three clusters, then a finer one of 5 groups. The partition *clust3* segregates the cells into three groups: Basal-like (c31 and c32) and Luminal (c33). Graphical results are in Figure 1.

**Fig. 1.**
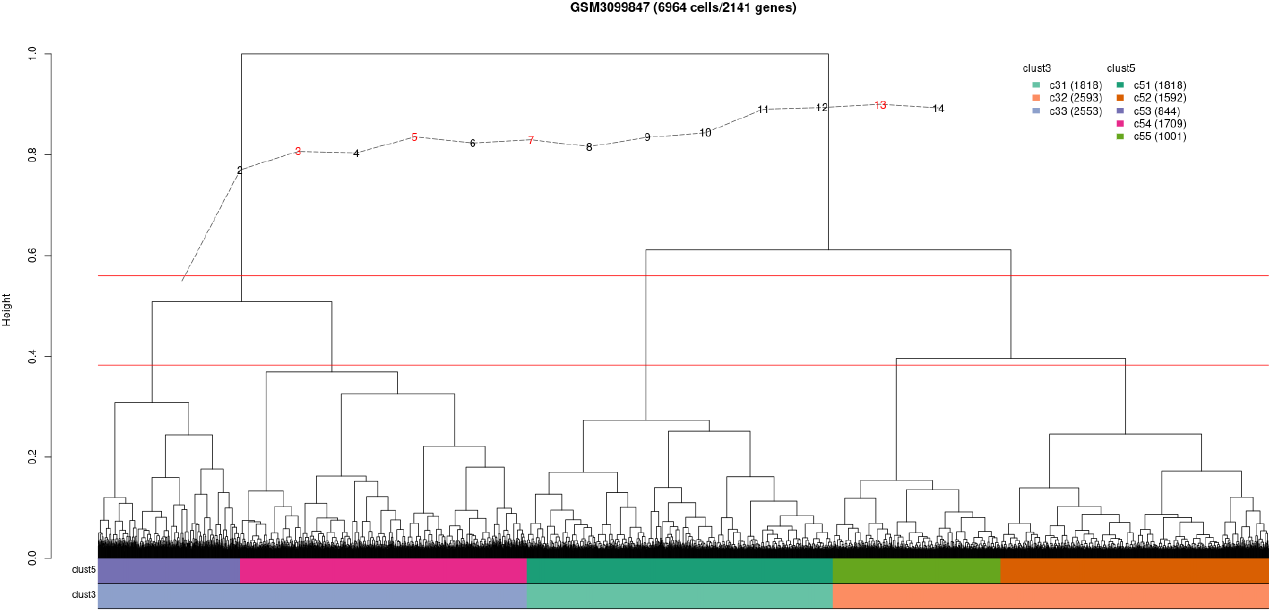
Hierarchical clustering with Ward’s linkage and Euclidean distance of 6,964 single-cell profiles and 2,141 features selected with AWST. The first annotation bar is the partition of cells into five groups (clust5); the second bar traces a partition of three clusters (clust3).

A sampling algorithm has been designed to detect the genes associated with each cluster. A synthetic control group was assembled by sampling the same number of items from each cluster; then, a gene-wise ANOVA was run to compare the synthetic control versus the clusters c31, c32, and c33, seen as treatment groups.

Figure 2a shows the statistical evidence of genes (top 200 features) uniquely associated with each cluster. Red means that aggregated gene level in the cluster is above average (synthetic control); green indicates that the level is below average. The coloring follows the log fold-change of each group to the synthetic control. With the help of signatures from [8], c32 is a population of basal-like/stem cells, and c33 is a luminal group. From [1], c31 is a proliferative set of cells.

**Fig. 2.**
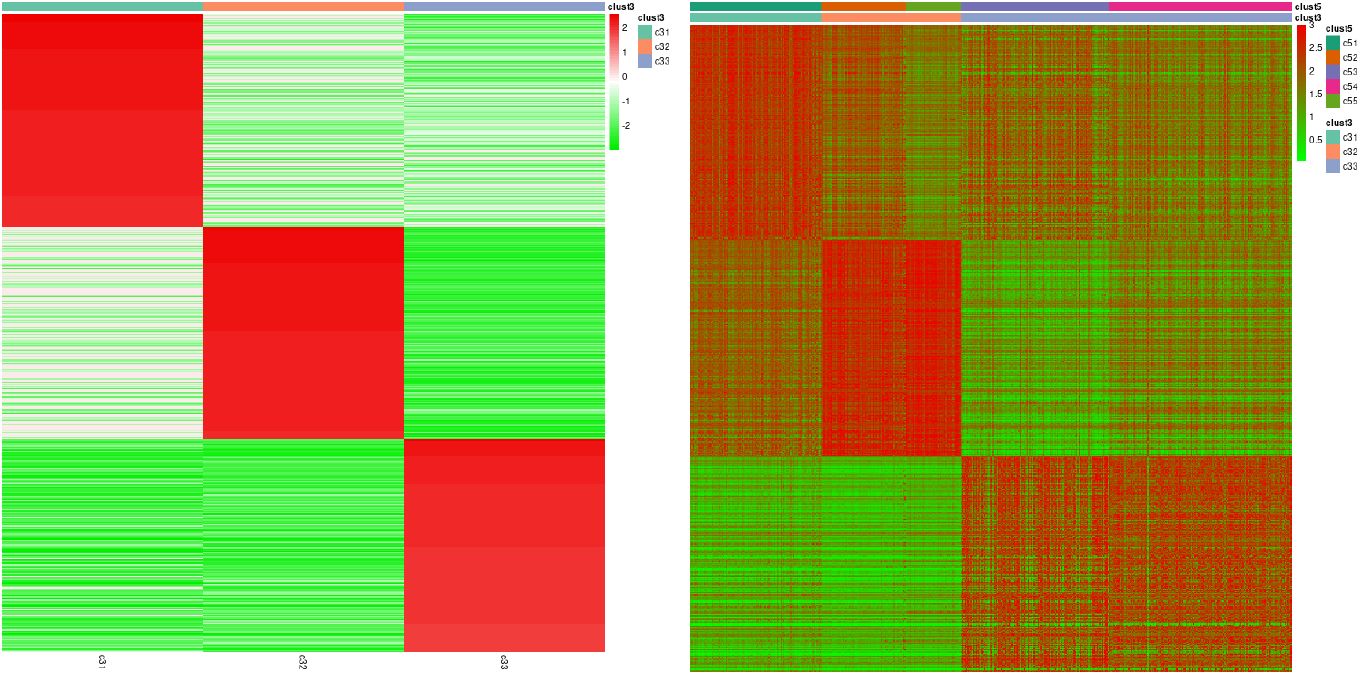
Study of clust3. Heat-map (left) of each cluster’s top 200 standardized log fold-changes in clust3 to a synthetic background. Green means under-regulation to the background level; red means up-regulation to the background. On the right: Heat-map of the top 200 gene expression levels associated with each cluster in clust3. Green means no expression level; red means high expression level.

Figure 2b shows the same genes as Figure 2a from a different point of view. This time the normalized gene levels have been rendered. Green means no expression. As the color moves to red, the expression level increases.

Figures 2a and 2b carry different information. The analysis of log fold-change to a common background suggests a trending behavior of cell population (more or less proliferative than the average); the study of expression levels provides an absolute trait of cells.

Figures 3a and 3b follow the same analytical process as before, but analyze clust5. The existence of sub-populations of cells in each of the significant aggregations in the three groups is evident.

**Fig. 3.**
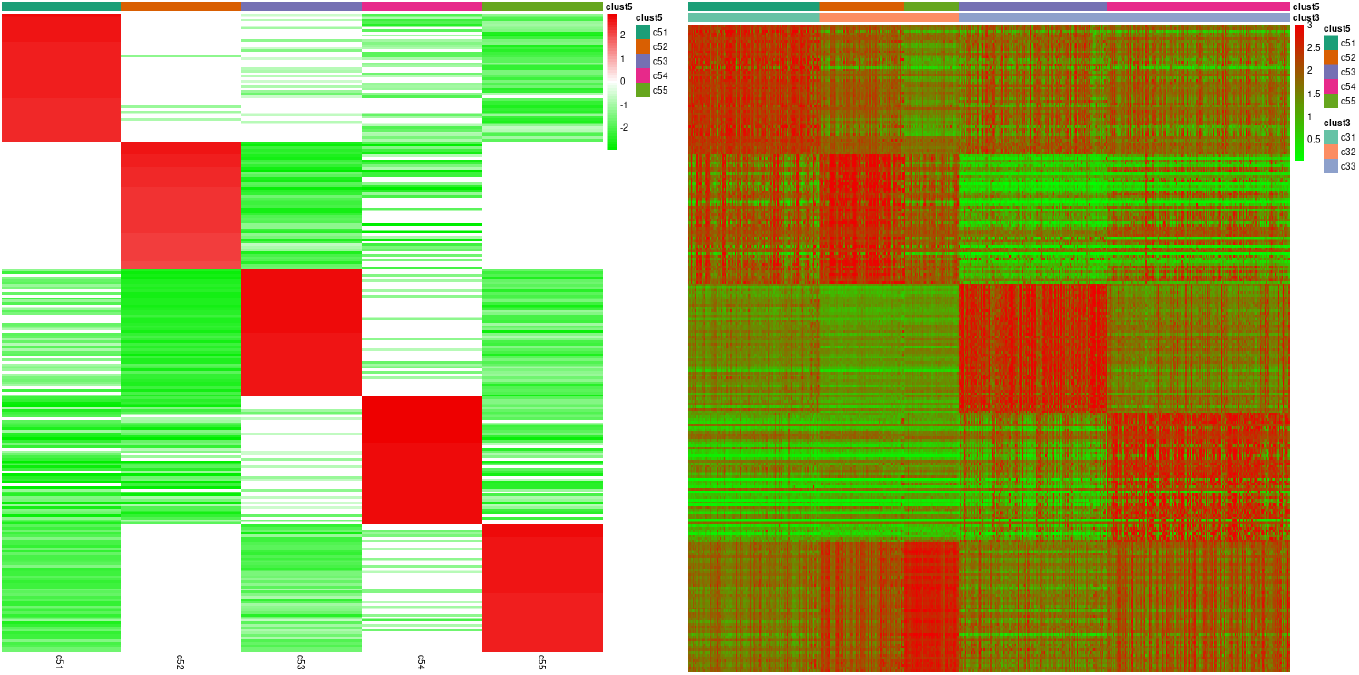
Study of clust5. Heat-map (left) of each cluster’s top 50 standardized log fold-changes in clust5 to a synthetic background. Green means under-regulation to the background level; red means up-regulation to the background. On the right: Heat-map of the top 50 gene expression levels associated with each cluster in clust5. Green means no expression level; red means high expression level.

## 3 Material

### Data

The example data are single-cells RNAseq of human breast epithelial cells analyzed in [11] and available with accession number GSM3099847 [6] at NCBI.

### Hardware

Superdome server (224 CPU cores Intel(R) Xeon(R) Platinum 8280 CPU @ 2.70GHz, 1.5 TB RAM) matched with docker.io/rocker/tidyverse:4.2.1

### Computational software

The analysis has been implemented in R version 4.2.3 (2023-03-15) – “Shortstop Beagle” (x86_64).

The program includes many libraries specified chunk by chunk. The software is available at [12] to facilitate reproducibility.

## 4 Methods and implementation

Here we show the essential steps. Tedious or routinely steps, such as data downloading, are not presented. They are in the script available from [12].

### Chunk n.1: downloading data

Retrieve the data (stored in `ddata` as a matrix) from the NCBI repository and save it in a local file `GSM3099847.RData`.

### Chunk n.2: pre-processing data

Apply the SAVER methodology to raw RNAseq data to recover gene expression [4, 5]

~~~
rm(list = ls())
load(“GSM3099847.RData”)
library(SAVER)
ddata <- saver(as.data.frame(ddata), estimates.only = TRUE, ncores = 60)
~~~

Then, a quantile normalization [13, 15] is applied to ddata

~~~
library(EDASeq)
ddata <- betweenLaneNormalization(ddata, which = “full”, round = FALSE)
save(ddata, file = “GSM3099847_SAVER_EDASeq.RData”)
~~~

This version of `ddata` is stored in `GSM3099847_SAVER_EdaSeq.RData`. This version of data is the base for any any downstream analysis.

### Chunk n.3: preparing data for clustering

Here the Asymmetric Within-Sample Transformation [**AWST_bioc**, 14] is applied to data, together with the gene-filtering. Then the usual Euclidean distance matrix is computed, followed by application of the hierarchical clustering with Ward’s linkage method.

~~~
rm(list = ls())
load(“GSM3099847_SAVER_EDASeq.RData”)
library(awst)
exprData <- awst(ddata, poscount = TRUE, full_quantile = TRUE)
dim(exprData <- gene_filter(exprData)) #[1] 2141 6964
exprData <- t(exprData)
nrow_exprData <- nrow(exprData)
ncol_exprData <- ncol(exprData)
ddist <- dist(exprData)
hhc <- hclust(ddist, method = “ward.D2”)
~~~

From the hierarchical data-structure the Calinski-Harabasz index is computed.

~~~
# full path in the .RMD file
source(“https://&/calinski_20210214.R“)
aCalinski <- calinski(hhc)
save(hhc, nrow_exprData, ncol_exprData, aCalinski,
   file = “GSM3099847_SAVER_EDASeq_AWST_hclust.RData”)
~~~

**Figure 1: Unsupervised hierarchical clustering**.

This chuck generates the dendrogram in Figure 1 and two data structures for later use. `clustering.df` is a data frame that collects the clusterings; `clust.colorCode` is instead a named list storing the association of each cluster with the corresponding color code. The variable `no_of_clusters` controls the cut of the dendrogram so that the required number of groups is gotten. From the set-up of this variable, automatic cluster association, naming, and color coding follow. Here, we show the essential steps. The full code is in [12].

~~~
clustering.prefix <- “clust”; short.prefix <- “c”
clustering.df <- data.frame(row.names = hhc$labels, barcode = hhc$labels)
clust.colorCode <- NULL
###
mmain <- paste0(“GSM3099847 (“, nrow_exprData, “cells/”, ncol_exprData, “genes)”)
hhc$height <- hhc$height/max(hhc$height)
plot(hhc, hang = -1, labels = FALSE, xlab = ““, sub = ““, main = mmain)
plot(aCalinski, add = TRUE, from = 500, to = 5000, shift = 0.55, height = 0.35, max_height = 1) ###
# this clustering is for later use
clustering.df$reduced <- as.factor(cutree(hhc, k = 500))
###
no_of_clusters <- 3
clustering_name <- paste0(clustering.prefix, no_of_clusters)
clustering_name.col <- paste0(clustering_name, “.col”)
hh <- (hhc$height[length(hhc$height)-no_of_clusters+2] +
     hhc$height[length(hhc$height)-no_of_clusters+1])/2
segments(1, hh, nrow_exprData, hh, col = “red”) # cut the tree
tmp <- as.factor(cutree(hhc, k = no_of_clusters)) # get the clusters
levels(tmp) <- paste0(short.prefix, no_of_clusters, c(paste(1:9), letters))[1:no_of_clusters] assign(clustering_name, tmp)
levels(tmp) <- brewer.pal(n = no_of_clusters, name = “Set2”)
assign(clustering_name.col, tmp)
clustering.df$tmp <- get(clustering_name)
clustering.df$tmp.col <- get(clustering_name.col)
tt <- table(clustering.df$tmp, clustering.df$tmp.col)
colorCode <- colnames(tt)[apply(tt, 1, which.max)]
names(colorCode) <- rownames(tt)
clust.colorCode <- c(clust.colorCode, colorCode)
tmp <- grep(“tmp”, colnames(clustering.df))
colnames(clustering.df)[tmp] <- gsub(“tmp”, clustering_name, colnames(clustering.df)[tmp])
~~~

After the running of this chunk, the content of `clustering.df` is

~~~
> head(clustering.df)
~~~

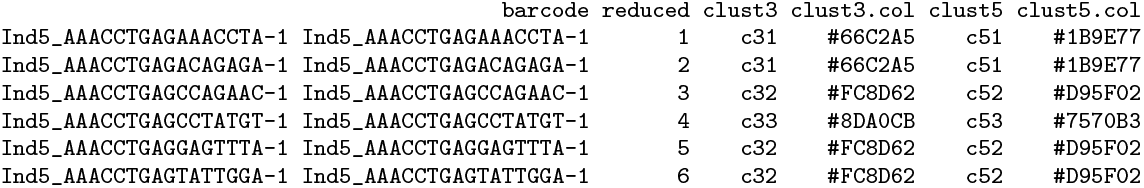

and the content of clust.colorCode is

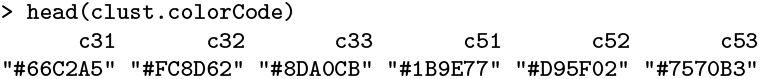

### Chunk 4: preparing data for Figures 2a and 2b

Figure 2 shows the result of a simulation experiment to discover the features associated with each cluster in the hierarchical clustering in Figure 1. In this chunk, we prepare data for the graphical display of heat maps.

The numerical results come from *B* = 50 montecarlo experiments where, each time, a synthetic background has been assembled. If we consider the clustering clust3, the synthetic control `c00` is a group of randomly sub-sampled (without replacement) cells from each of the other clusters c31, c32, and c33 in equal numbers. This way, a new clustering clust3_*b*_ rises, with one more cluster than the original partition, *b* = 1, 2, …, 50. clust3_*b*_ still classifies the *n* = 6,964 initial cells. This is an example of a random clust3_*b*_ (`clust3` is the original partition, and `design` corresponds to clust3_*b*_).

~~~
> addmargins(table(annotation.tmp$design, annotation.tmp$clust3))
~~~

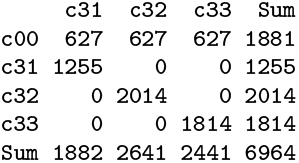

From each of the clusters in clust3_*b*_, a sample with replacement (like bootstrap) of dimension *m* is sampled. *m* is set to one-third of the minimum size of the clusters in clust3_*b*_. clust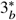 is the collection of the “*bootstrap*” clusters. This an example of the bootstrap partition

~~~
>addmargins(table(annotation.tmp$design, annotation.tmp$clust3))
~~~

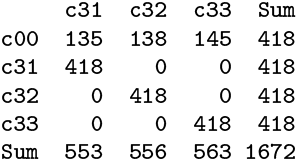

Clust 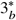 transformed in a model matrix **X**, feeds the DESeq2 [10, 9] procedure to estimate a generalized linear model. In each row of **X**, all elements are zero but one set to 1. Given a gene j,

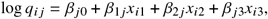

where *x*_*i*1_, *x*_*i*2_, and *x*_*i*3_ are elements of the *i*^*th*^ row of **X**, and *q*_*i j*_ is proportional to the true expected counts of the *j* ^*th*^ in the *i*^*th*^ samples [10, 2]. The intercept estimate 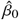 is the average of the *j* ^*th*^ gene levels in `c00`. Given the setup, the estimate of *β*_*k j*_, *k*> 0, is the difference between the average of levels in the *k*^*th*^ cluster and the average in `c00`. Essentially, 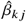 is a fold-change estimate between the gene level in the *k*^*th*^ cluster and the reference cluster `c00`.

From each *b* run, we retain Wald’s statistics associated with each *β*’s. Wald’s statistic is a standardized log_2_ fold-change with an asymptotic Gaussian distribution.

The construction of the partitions clust3_*b*_ and clust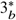, the setup and the running of the linear model with DESeq2 are all enclosed in the function `single_shot()`. The value of such a function is the output of `DESeq()` function.

~~~
single_shot <- function() {
annotation.tmp <- clustering.df
mmin <- min(tt <- table(annotation.tmp$design))
nooc <- length(tt) # no of clusters
# assemble clust3$_b$
tmp <- c()
for(this_cluster in unique(annotation.tmp$design)) tmp <- c(tmp,
     sample(annotation.tmp$barcode[annotation.tmp$design == this_cluster]
     mmin/nooc, replace = FALSE))
annotation.tmp$design[annotation.tmp$barcode %in% tmp] <-“000” # synthetic background
# assemble clust3$_b^*$
m <- min(table(annotation.tmp$design))/3
tmp <- c()
for(this_cluster in unique(annotation.tmp$design))
tmp <- c(tmp,
    sample(annotation.tmp$barcode[annotation.tmp$design == this_cluster], m, replace = TRUE)) annotation.tmp <- annotation.tmp[tmp,]
# rename the items
for(i in 1:nrow(annotation.tmp))
   annotation.tmp$id[i] <- paste(sample(c(letters, LETTERS, 0:9), 15, replace = TRUE), collapse = ““)
# setup the data-structure for DESeq2
data.tmp <- ddata[, annotation.tmp$barcode]
colnames(data.tmp) <- annotation.tmp$id
rownames(annotation.tmp) <- annotation.tmp$id
annotation.tmp$design <- factor(annotation.tmp$design)
dds <- DESeqDataSetFromMatrix(countData = data.tmp, colData = annotation.tmp, design = ∼ design)
# in DESeq2, suggested correction for single cell data
tmp <- scran::computeSumFactors(dds)
dds@colData@listData[[“sizeFactor”]] <- tmp@colData@listData$sizeFactor
ans_DESeq <- DESeq(dds, test = “Wald”, useT = T,
   minmu = 1e-6, fitType=‘local’, minReplicatesForReplace = Inf)
ans_DESeq <- mcols(ans_DESeq, use.names=TRUE)
ans_DESeq <- data.frame(ans_DESeq) invisible(ans_DESeq)
}
~~~

*B* = 50 times the `single_shot()` is called; Wald’s statistics are extracted from the output and averaged across the runs. The strong law of large numbers assures that the average values be good approximations of the true values of the log_2_ fold-change (Wald’s statistics) of each genes for each cluster to a generic background.

The result of this chunk is stored for the visualization task.

~~~
rm(list = ls())
load(“GSM3099847_SAVER_EDASeq.RData”)
# makes expression values similar to counts per millions (CPM)
ddata <- apply(ddata, 2, function(x) x/sum(x)*1e6)
mode(ddata) <- “integer” # transforms in integer with an implicit truncation
load(“GSM3099847_SAVER_EDASeq_AWST_hclust_clustering.RData”)
clustering.df$design <- as.character(clustering.df$clust3)
clustering.df$id <- NA
library(DESeq2)
library(scran)
ans <- single_shot()
WaldStatistic_design <- grep(“WaldStatistic_design”, colnames(ans)
tmp <- ans[, WaldStatistic_design]
runs <- 50
k <- 2
while(k <= runs) {
system.time(ans <- single_shot())
tmp <- tmp + ans[, WaldStatistic_design]
k <- k + 1
}
ans <- tmp/runs
ans <- ans[-which(is.na(ans[, 1])),]
colnames(ans) <- unique(clustering.df$design)
save(ans, file = “GSM3099847_SAVER_EDASeq_ansDESeq2_clust3.RData”)
~~~

**Figure 2a: Heat-map of each cluster’s top 200 standardized log fold-changes in clust3 to a synthetic background**.

Figure 2 shows the result of a simulation experiment to discover the features associated with each cluster in the hierarchical clustering in Figure 1.

This chunk analyzes preparatory data from the montecarlo experiment and adapts to be displayed with the `pheatmap` [7] function. Here, we comment just some relevant and ‘obscure’ step.

~~~
rm(list = ls())
load(“GSM3099847_SAVER_EDASeq_AWST_hclust_clustering.RData”)
load(“GSM3099847_SAVER_EDASeq_ansDESeq2_clust3.RData”)
colnames(ans) <- c(“c31”, “c32”, “c33”)
> head(ans)
~~~

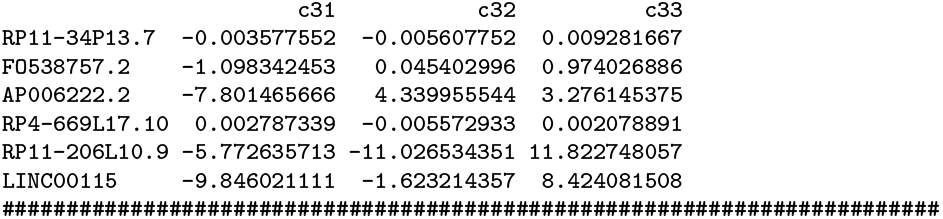

`ans` is the data-frame where each column is the Wald’s test statistics associated with each cluster and for each gene (row-wise). From the test value, we get the *p*-values and adjust for multiple tests, according to Benjamini-Hochberg’s correction.

~~~
p.values <- 2*(1-pnorm(abs(as.matrix(ans))))
p.values <- apply(p.values, 2, p.adjust, method = “BH”)
~~~

Given a level of significance `alpha`, we set to zero every non-significant entry of `ans`; then, we restrict the data frame to rows with only one component greater than zero.

~~~
alpha <- 0.01
ans <- ans * (p.values < alpha)
ans <- ans[rowSums(ans > 0) == 1,]
~~~

A statement like `ans[rowSums(ans > 0) == 2,]` would have selected the rows having two elements greater than zero. Then the selected genes are up-regulated in two clusters. Having many clusters, the condition allows the selection of genes up-regulated in any combination of groups.

~~~
max_tops <- 200
ttable <- table(colnames(ans)[apply(ans, 1, which.max)])
K <- sapply(ttable, function(x) min(max_tops, x))
j <- 1
tops <- head(rownames(ans[order(ans[, j], decreasing = TRUE),]), K[j])
for(j in 2:ncol(ans))
tops <- c(tops, head(rownames(ans[order(ans[, j], decreasing = TRUE),]), K[j]))
~~~

The following line implements a scaling of the *w*_*i j*_ Wald’s test values to remove differences in variability associated with clusters:

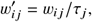

where *j* ^*th*^ is associated with clusters and *i*^*th*^ with genes. The *w*_*i j*_ ‘s are stored in the `ans` matrix. The scaling factor *τ*_*j*_ is a dispersion index from zero

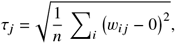

where *n* is the number of genes in `ans`. This way, the original zero value of data is preserved since it means that there is no fold-change of the level for a cluster to the baseline (synthetic cluster `c00`).

~~~
ans <- apply(ans[tops,], 2, function(x) x/sqrt(mean(x^2)))
~~~

The next step controls the contrast in the heat map and restricts the values in the range from -3 to +3. Values greater than 2*τ* from zero are full red; symmetrically, values lower than 2*τ* from zero are full green.

~~~
ans <- 6 * (pnorm(ans) - 0.5)
annotation_col <- data.frame(clust3 = colnames(ans), row.names = colnames(ans))
ann_colors = list(clust3 = clust.colorCode[grep(“c3”, names(clust.colorCode))])
suppressPackageStartupMessages(library(pheatmap))
pheatmap(ans, cluster_rows=FALSE, cluster_cols=FALSE, show_rownames = FALSE,
       annotation_col=annotation_col,
       annotation_colors = ann_colors,
       color = colorRampPalette(c(“green2”, “white”, “red2”))(51))
~~~

Finally, in the heat map of Figure 2 have been shown the levels of the standardized fold-change (Wald’s test value) associated with each cluster and with respect to the synthetic control group c00.

**Figure 2b: Heat-map of the top 200 gene expression levels associated with each cluster in clust3**.

This chunk has in charge of displaying the heat map of the expression level of the genes for each cell. Being the number of cells above the thousands, we aggregate the cells in 500 small clusters. Such an aggregation comes from chunk ‘Figure 2’, where we set

~~~
clustering.df$reduced <- as.factor(cutree(hhc, k = 500))
~~~

The level associated with each cluster is summarized as the average of the levels participating in the group. `pheatmap` allows a dimensionality reduction but follows algorithms that can corrupt the external ordering coming from the hierarchical clustering.

~~~
load(“GSM3099847_SAVER_EDASeq.RData”)
###
rdata <- as.data.frame(t(ddata[tops,]))
rdata <- aggregate(rdata, list(reduced = clustering.df$reduced), mean)[, -1]
~~~

In the following two steps, the same transformation as in the chunk ‘Figure 2’ have been applied to data before feeding `pheatmap`. The meaning of the color is now different from Figure 2. The gene level’s zero value (in green) means no expression (negative intensities are not allowed). The gene levels are in red in the case of up-regulation.

~~~
rdata <- apply(rdata, 2, function(x) x/sqrt(mean(x^2)))
rdata <- t(rdata)
colnames(rdata) <- paste0(“R”, 1:ncol(rdata))
rdata <- 6*(pnorm(rdata) - 0.5)
tt <- table(clustering.df$reduced, clustering.df$clust3)
tt <- colnames(tt)[apply(tt, 1, which.max)]
annotation_col <- data.frame(row.names = colnames(rdata), clust3 = tt)
tt <- table(clustering.df$reduced, clustering.df$clust8)
tt <- colnames(tt)[apply(tt, 1, which.max)]
annotation_col$clust8 <- tt
annotation_col <- annotation_col[order(annotation_col$clust3, annotation_col$clust8),]
rdata <- rdata[, rownames(annotation_col)]
ann_colors <- list(clust3 = clust.colorCode[grep(“c3”, names(clust.colorCode))],
                 clust8 = clust.colorCode[grep(“c8”, names(clust.colorCode))])
pheatmap(rdata, cluster_rows=FALSE, cluster_cols=FALSE,
       show_colnames = FALSE, show_rownames = FALSE,
       annotation_col = annotation_col,
       annotation_colors = ann_colors,
       color = colorRampPalette(c(“green”, “red2”))(50))
~~~

